# Sex-specific Regulation of Fentanyl Reward by the Circadian Transcription Factor NPAS2

**DOI:** 10.1101/2024.11.12.623242

**Authors:** Kelly Barko, Micah A. Shelton, Lauren M. DePoy, Jenesis Gayden-Kozel, Sam-Moon Kim, Stephanie Puig, Xiangning Xue, Puja K. Parekh, George C. Tseng, Benjamin R. Williams, Jeffrey Oliver-Smith, Xiyu Zhu, Zachary Freyberg, Ryan W. Logan

**Affiliations:** Department of Psychiatry, University of Pittsburgh School of Medicine, Pittsburgh, PA, USA; Department of Psychiatry, University of Massachusetts Chan Medical School, Worcester, MA, USA; Department of Neurobiology, University of Massachusetts Chan Medical School, Worcester, MA, USA; Department of Biostatistics, University of Pittsburgh, Pittsburgh, PA, USA; Department of Neuroscience, University of Texas at Dallas, Dallas, TX, USA; Department of Cell Biology, University of Pittsburgh, Pittsburgh, PA, USA

## Abstract

Synthetic opioids like fentanyl are highly potent and prevalent in the illicit drug market, leading to tolerance, dependence, and opioid use disorder (OUD). Chronic opioid use disrupts sleep and circadian rhythms, which persist even during treatment and abstinence, increasing the risk of relapse. The body’s molecular clock, regulated by transcriptional and translational feedback loops, controls various physiological processes, including the expression of endogenous opioids and their receptors. The circadian transcription factor NPAS2, highly expressed in the nucleus accumbens, may have a crucial function in opioid-related behaviors. Our study found sex-specific roles for NPAS2-mediated reward behaviors in male and female mice, including in fentanyl seeking and craving. We also identified specific cell types and transcriptional targets in the nucleus accumbens of both mice and humans by which NPAS2 may mediate the impact of fentanyl on brain physiology and in opioid reward-related behaviors. Ultimately, our findings begin to uncover the mechanisms underlying circadian rhythm dysfunction and opioid addiction.

## INTRODUCTION

Synthetic opioids, such as fentanyl, are highly potent, addictive substances that dominate the illicit drug supply. Repeated use of fentanyl can lead to tolerance and dependence, contributing to the risk of developing opioid use disorder (OUD). Common consequences of chronic opioid use and misuse are sleep and circadian rhythm disruptions, often persisting during treatment and abstinence^1^. Moreover, sleep and circadian alterations contribute to the overall risk of relapse to opioids and other substances^2^. Further studies to investigate these links between circadian pathways and opioid addiction are clearly needed and could eventually lead to novel therapeutics for OUD.

Nearly every cell in the body contains a molecular clock, controlled by a series of transcriptional and translational feedback loops^3^. Sets of transcription factors drive the expression of clock-controlled genes (CCGs) in a tissue-dependent and cell type-specific manner to regulate specific metabolic, physiological, and signaling pathways^4^. In peripheral tissues, such as spleen, and the brain, the molecular clock controls the expression of endogenous opioids and opioid receptors. The circadian transcription factor Npas2 is predominantly expressed in the spinal cord and brain, enriched in several brain regions involved in reward, motivation, and addiction^5,6^. Prior work has shown that the expression and transcriptional activity of Npas2 is comparatively higher and preferentially enriched in the nucleus accumbens of the striatum^6,7^—a key neural hub in the regulation of opioid-related behaviors. Yet, important gaps in our knowledge exist, particularly regarding whether: 1) opioids like fentanyl alter Npas2 expression in the nucleus accumbens, 2) if some striatal cell types are more affected than others by opioid actions, and 3) if these opioid-induced expression changes are reflected at the behavioral level.

In our current work, we investigated the role of Npas2 in opioid reward-related behaviors, including opioid seeking, craving, and relapse behaviors in male and female mice. We used global NPAS2-deficient mice which carry a LacZ reporter replacing exon 2 that effectively excises the basic Helix-Loop-Helix (bHLH) domain. The bHLH domain is necessary for the transcription factor NPAS2 to directly bind DNA, and thus, prevents NPAS2-dependent circadian transcription creating NPAS2-deficient mice. We narrowed the potential actions of NPAS2 to regulate opioid-related behaviors to specific cell types in the nucleus accumbens, while also identifying cell type-specific transcriptional targets of NPAS2 associated with opioid-induced behavioral changes and human OUD.

## METHODS

### Mice

Mating of heterozygous NPAS2-deficient mice^6^ reproduced wild-type littermates with homozygous mutants, NPAS2-deficient mice, maintained on a C57Bl/6J background. Male and female wild-type and NPAS2-deficient mice were used for experiments (12-18 weeks, n=8-12/sex/group for behavioral experiments and n=6-8/sex/group for gene expression and RNA-sequencing experiments). NPAS2-deficient mice have the coding region of the bHLH domain replaced with a modified β-galactosidase reporter construct (LacZ). Mice were group housed and maintained on 12:12 light-dark cycle lights on at 700 and off at 1900, corresponding to Zeitgeber time (ZT) 0 and 12, respectively. Standard laboratory chow and water were provided *ad libitum*. Animal use was conducted in accordance with both ARRIVE^8^ and National Institute of Health guidelines and approved by the Institutional Animal Care and Use Committees at the University of Pittsburgh, Boston University School of Medicine, and University of Massachusetts Chan Medical School.

### Drug

Fentanyl hydrochloride was procured from National Institute on Drug Abuse drug supply.

### Fentanyl Conditioned Place Preference

Mice were assayed for conditioned drug reward using conditioned place preference (CPP)^7^. Briefly, mice were habituated to the testing room for 30 minutes prior to daily testing and conditioning. On pre-test day 1, mice freely explored the three-compartment place-conditioning apparatus. Time spent in each of the three compartments was measured and preference for any of the compartments was assessed. Most of the mice (>75%) exhibited a bias towards one of the conditioning compartments or the other. Thus, we used a biased conditioned protocol during which mice were conditioned on the less preferred compartment. On days 1, 3, and 5, mice were injected with saline (i.p.; 10ml/kg), then placed into one of the preferred compartments for 20 minutes. On days 2, 4, and 6, mice were administered fentanyl (i.p.; 0.1, 0.5, or 1mg/kg), then placed into the less preferred compartment for 20 minutes. On day 7, mice freely explored each of the three compartments of the conditioning apparatus. Time spent in each compartment was measured. The score for CPP was calculated by subtracting the time spent on the saline-paired compartment from the fentanyl-paired compartment. Mice were assayed between ZTs 3-5.

### Fentanyl Locomotor Sensitization

Mice were administered saline for two consecutive days to establish injection altered baseline activity. Following saline, mice were administered fentanyl (i.p.; 0.1mg/kg) for five consecutive days. Locomotor activity was tracked following injections in daily 1-hour sessions in an open field with infrared beam tracking (Kinder Scientific, Poway, CA). On each day of saline or fentanyl administration, mice were placed temporarily (~10 minutes) in a clean holding cage before being placed into the open field. On days 8-10, mice received no injections, followed by a challenge day (day 11), during which mice were administered fentanyl (i.p.; 0.1mg/kg). Locomotor activity was tracked using MotorMonitor software (Kinder Scientific).

### Fentanyl Oral Self-Administration

For operant self-administration, we used extra-wide modular mouse test chambers (ENV-307W-CT; Med-Associates, Fairfax, VT). Each chamber was housed within a larger sound-attenuating cabinet, equipped with a house light, stimulus lights, and ultra-sensitive retractable mouse levels, with dual pellet/liquid dipper dispensers and head-entry detectors. The dual pellet/liquid dispenser was located centrally in a port between the levers and stimulus lights were located directly above each lever. We specifically chose an operant self-administration assay of oral fentanyl consumption^9,10^, since oral routes of fentanyl administration are highly prevalent^11^. Prior to fentanyl self-administration, mice were trained on oral self-administration of sucrose.

Mice underwent 2-hour daily sessions of sucrose self-administration. Mice received food and water *ad libitum*, except for a 12-hour period immediately prior to the start of sucrose training, during which only water was restricted. Active lever presses raised the liquid dispenser arm that was equipped with a 100 μl dipper containing sucrose (10% w/v). The dipper remained in the raised position until the mouse completed a head entry. During this period, the stimulus light located directly above the active lever was lit and the house light turned off. After head entry (10 seconds), the dipper retracted and the house light re-activated. Inactive lever presses produced no activation of dipper, stimulus light, or house light. Sucrose training occurred from training days 1-7, depending on when criteria thresholds were consistently met (≥ 50 rewards for three consecutive days).

Once mice passed criteria, they were transitioned to oral fentanyl self-administration. The assignment of active versus inactive levers were switched relative to sucrose self-administration to prevent response bias during the initial sessions of fentanyl self-administration. An active lever press delivered 100 μl of fentanyl-containing solution of varying doses depending on acquisition and dose-response sessions. Delivery of fentanyl was accompanied by a 10 second tone and stimulus light. Inactive lever presses resulted in no change. During each daily session, the amount of solution consumed, number of active and inactive lever presses, were recorded. Each daily session was 2-hour in duration, except for progressive ratio sessions, during which sessions were 4 hours. During *acquisition* of fentanyl self-administration, mice received 1 μg/ml of fentanyl solution based on a fixed-ratio 1 (FR1) schedule.

During acquisition, mice received a 1 μg/ml reward on a FR1 schedule. Daily sessions were performed once per day for a minimum of 10 days until criteria was met (≥50 rewards and ≥66% ratio of active lever presses/total lever presses). Following acquisition, mice underwent a *descending dose response* (10, 3, 1, 0.3, 0.1, 0 μg/ml), during which each dose was presented for two days (12 days overall). Mice were then required to self-administer fentanyl for two days at a baseline dose of 1ug/ml to meet acquisition criteria again for at least one day before proceeding to progressive ratio. During *progressive ratio*, mice self-administered three doses of fentanyl (3, 1, 0.3 μg/ml) for two days per dose (6 days overall). Mice were again required to administer for a minimum of two days to meet acquisition criteria for at least one day before proceeding to extinction. During *extinction*, any press at the active lever was absent of any reward, tone, stimulus or house lights. Extinction lasted for 10 consecutive days of daily sessions. Following extinction, mice were tested for *cue-induced reinstatement* behavior for two days, during which an active lever press produce the tone, stimulus light, activation of the dipper, in the absence of the fentanyl reward. Behavioral responses were recorded using MED-PC IV software (Med-Associates). The volume of fentanyl solution consumed during each daily session was recorded.

### RNA Isolation and Gene Expression Assays

Mice were administered either saline, acute (one day) or chronic fentanyl (seven days; i.p.; 10ml/kg; 1mg/kg), then sacrificed 24 hours following last injection using rapid decapitation. Tissue homogenization was followed by 15-mins of treatment with DNAse I to digest any remaining genomic DNA, according to manufacturer’s protocols (Invitrogen). Total RNA (1ug) was used to synthesize cDNA with Superscript III Reverse Transcriptase (Invitrogen). cDNA was mixed with SYBR Green master mix (Applied Biosystems, ABI) and specific primers or probes (ABI) for genes or promoter regions of interest in mice (*Npas2* F-5’-GACACTGGAGTCCAGACGCAA, R-5’-AATGTATACAGGGTGCGCCAAA; *Clock*: F-5’-CAGAACAGTACCCAGAGTGCT, R-5’-CACCACCTGACCCATAAGCAT; *B2m* F-5’-GGGTGGAACTGTGTTACGTAG, R-5’-TGGTCT TTCTGGTGCTTGTC). Prior to the experiment, primer sets were tested thoroughly to determine reaction efficiency, specificity, and the absence of primer-dimers. Reactions were run on an ABI Prism 7700 real-time PCR machine. Fold changes and relative gene expression were calculated using the comparative Ct method and normalized to the corresponding *Gapdh* or *18S* mRNA levels.

### Cell Type-Specific Isolation and RNA-Sequencing

To sequence activity-translated mRNAs, male and female *Drd1a-Cre* mice were crossed to RiboTag mice expressing a Cre-inducible HA-Rpl22^12^. Male and female mice were between the ages of 12-14 weeks and maintained on 12:12 light-dark cycle (Lights on at 700). Mice were administered fentanyl (i.p.; 1mg/kg) for seven consecutive days then sacrificed by rapid decapitation 24-hrs following the last injection at either ZT6 or ZT18. Nucleus accumbens tissue was collected from male and female mice across two times of day (ZT6 and ZT18). Ribosomal mRNAs from cells expressing *Drd1* were isolated via co-immunoprecipitation^12^. Libraries were prepared and sequenced with experimenters blinded to groups. Libraries were sequenced (NextSeq 500 and Illumina). After alignments (*Mus musculus* Ensembl GRCm38; HISAT2), preprocessing and filtering of sequencing data, low expression transcripts were filtered, retaining transcripts with at least one count per million (CPM) in half of the samples. After filtering, differential expression (DE) was analyzed using DESeq2^13^ with main and interaction effects of sex, drug, and ZT. Pathway enrichment analyses were conducted using Metascape.org^1^. Using diurnal rhythm NPAS2 chromatin immunoprecipitation (ChIP) datasets^7^, we compared NPAS2 bound genes to DE genes at ZT6 and ZT18, focusing only on promoter sequences, at the interaction of sex and condition (saline versus fentanyl).

### Stereotaxic surgery

Stereotaxic surgery was performed as previously described^7^. Briefly, mice underwent stereotaxic brain surgery to bilaterally inject the nucleus accumbens (relative to bregma: angle 10°; AP, ±1.5; ML, ±1.5; DV, −4.4). High-titer viruses (1 μl) encoding either AAV2/2.H1lox.mCherry-Npas2-(or Scramble)-shRNA^2^ was injected into the nucleus accumbens of male or female *Drd1a*-Cre and *Drd2*-Cre mice (12-14 weeks of age). Following stereotaxic surgery, mice recovered for ~3 weeks prior to behavioral testing to allow for maximal viral expression and *Npas2* knockdown. Mice then underwent fentanyl CPP, as described above. shRNA-mediated knockdown was validated using micro-dissected nucleus accumbens punches from mice following maximal viral expression. Coronal sections containing the nucleus accumbens with mCherry reporter of AAV expression were visualized using NightSea BlueStar flashlight under a dissecting microscope. Punches with concentrated mCherry expression were dissected from nucleus acccumbens sections for gene expression assays to verify knockdown of *Npas2* expression.

### Multiplex RNAscope fluorescent *in situ* hybridization

#### RNAscope

Multiplex RNAscope technology (ACD Bio, Neward, CA) measured mRNA expression of *Drd1*, *Drd2*, and *Npas2* genes. RNAscope labelling of fresh frozen brain tissue was conducted using the ACD Bio RNAscope v1 kit as described earlier^14–16^. Briefly, pre-mounted tissue sections were post-fixed on slides in pre-chilled 4% paraformaldehyde (4°C, 60 minutes). Tissue underwent dehydration in successive ethanol baths of increasing concentration (50%, 70%, 100%). Upon drying, samples were incubated with probes (2 hours, 40°C) to detect cell-specific mRNA expression of mouse *Drd1* (ACD Bio, Mm-Drd1, Cat. 461901-C3), *Drd2* (Mm-Drd2, Cat. 406501), *and Npas2* (Mm-Npas2, Cat. 493961-C2). After hybridization, the respective signals were amplified with the following fluorophores: Channel 1, Atto 550; Channel 2, Alexa fluor 647; Channel 3, Alexa fluor 488. Tissues were counter-stained with DAPI to label cell nuclei and mounted with Vectashield (Vector Labs, Neward, CA). Slides were stored at 4°C until imaging.

#### Fluorescence slide scanning microscopy

Fluorescence imaging of labeled slides was performed using the Olympus VS120 automated slide scanner (Olympus, Center Valley, PA). Exposure times were adjusted for each fluorophore for accurate signal distribution with no signal saturation. Coronal sections were identified and imaged as 3μm-thick Z stacks (3 Z-planes spaced by 1μm each) using a dry 20X objective lens (N.A. 0.8). Identical exposure settings and magnifications were consistently applied to all slides. After image acquisition, Z stacks were converted to two-dimensional maximum intensity projections.

#### Image analysis

Image analysis was performed using the HALO image analysis platform equipped with a fluorescent *in situ* hybridization module (Indica Labs, Albuquerque, NM). Nuclei were quantified as DAPI-stained objects with the minimum cytoplasmic radius set at 5μm. Puncta corresponding to the respective mRNA probes were quantified as any 0.03-0.15μm^2^ object. We normalized mRNA grain numbers per probe by dividing by the number of grains by total area analyzed (mRNA grains/area analyzed). We then tested several thresholds based on our prior studies^17^. Based on these analyses, we employed a 5x threshold above the puncta/area analyzed values which was optimal for accurate identification of labelled cells. We calculated the cellular density of positive cells by dividing the number of positive cells by the total area analyzed and converted these values to mm^2^ units.

### Integration of Mouse and Human Postmortem Brain Datasets

Prior work from us identified alterations in transcript rhythms in postmortem human nucleus accumbens in individuals with OUD^18^. Using ChEA3^19^, a web-server application developed to conduct transcription factor enrichment analysis, we identified transcription factor networks predicted from the transcripts that lost rhythmicity in individuals with OUD^18^—406 transcripts. ChEA3 integrates data about transcription factor/target– gene associations from multiple assay types and other sources of evidence. Transcription factors are prioritized based on the overlap between user-inputted gene sets and annotated sets of transcription factor targets stored within the ChEA3 database^19^. ChEA3 identified a network of interacting transcription factors predicted from the lost of transcript rhythmicity in the human nucleus accumbens in OUD.

### Statistical Analyses

GraphPad Prism software (10.4.0, GraphPad Software, LLC) was used for statistical analyses. Differences in conditioned place preference measured as the time spent in the drug-paired context were analyzed using a two-way analysis of variance (ANOVA), genotype by dose. Locomotor sensitization, oral sucrose self-administration, and oral fentanyl self-administration were also analyzed by two-way ANOVA, with main effects of genotype and day, except for the dose response study, which used factors genotype and dose. Differences in gene expression following fentanyl administration were assessed using two-way ANOVA with treatment as main effect (saline, acute, or chronic). One-way ANOVA followed by Tukey’s multiple comparison tests were employed to analyze differences in the density of dopamine D1 and D2 receptor-expressing cells and *Npas2* mRNA expression across striatal subregions. Significant interactions from ANOVAs were further investigated using Tukey’s or Sidak’s multiple comparisons tests.

## RESULTS

### Increased fentanyl conditioned place preference and locomotor sensitization in male and female Npas2-deficient mice

To examine the role of Npas2 in opioid reward, we used conditioned place preference (CPP) behavioral assays with fentanyl administration. Both male and female NPAS2-deficient mice spent significantly more time in the fentanyl-paired chamber across each of the fentanyl doses (**Figure 1A**; genotype by dose interaction in males, F(2, 42) = 4.7, p=0.014 with Sidak’s multiple comparisons test identifying significant genotype differences at each dose, p<0.01; and main effect of genotype in females, F(1, 42) = 27, p<0.0001). Wild-type males also had increased fentanyl CPP at higher doses (**Figure 1A**). We further investigated the impact of Npas2 deficiency on fentanyl-induced locomotor sensitization, a behavior considered to be considered as associated with opioid-seeking and opioid-taking behaviors^20^. Repeated fentanyl administration led to significantly increased locomotor activity across successive days in both wild-type and Npas2-deficient males (**Figure 1C**; genotype by day interaction, F(7, 154) = 2.4, p=0.025 with Tukey’s multiple comparisons test identifying significant differences between genotypes on days 1, 3-10, p<0.05) and female mice (**Figure 1D**; main effects of day, F(3.4, 74.8) = 21.1, p<0.0001, and genotype, F(1, 22) = 8.6, p=0.008). We found no meaningful differences between day 7, the last day of repeated fentanyl injections, relative to day 10, “challenge” day (p>0.05). Overall, the loss of Npas2 function leads to increased fentanyl conditioned reward and sensitization in male and female mice, with a few sex-specific effects on fentanyl-related reward behaviors by dose and administration frequency.

**Figure 1.**
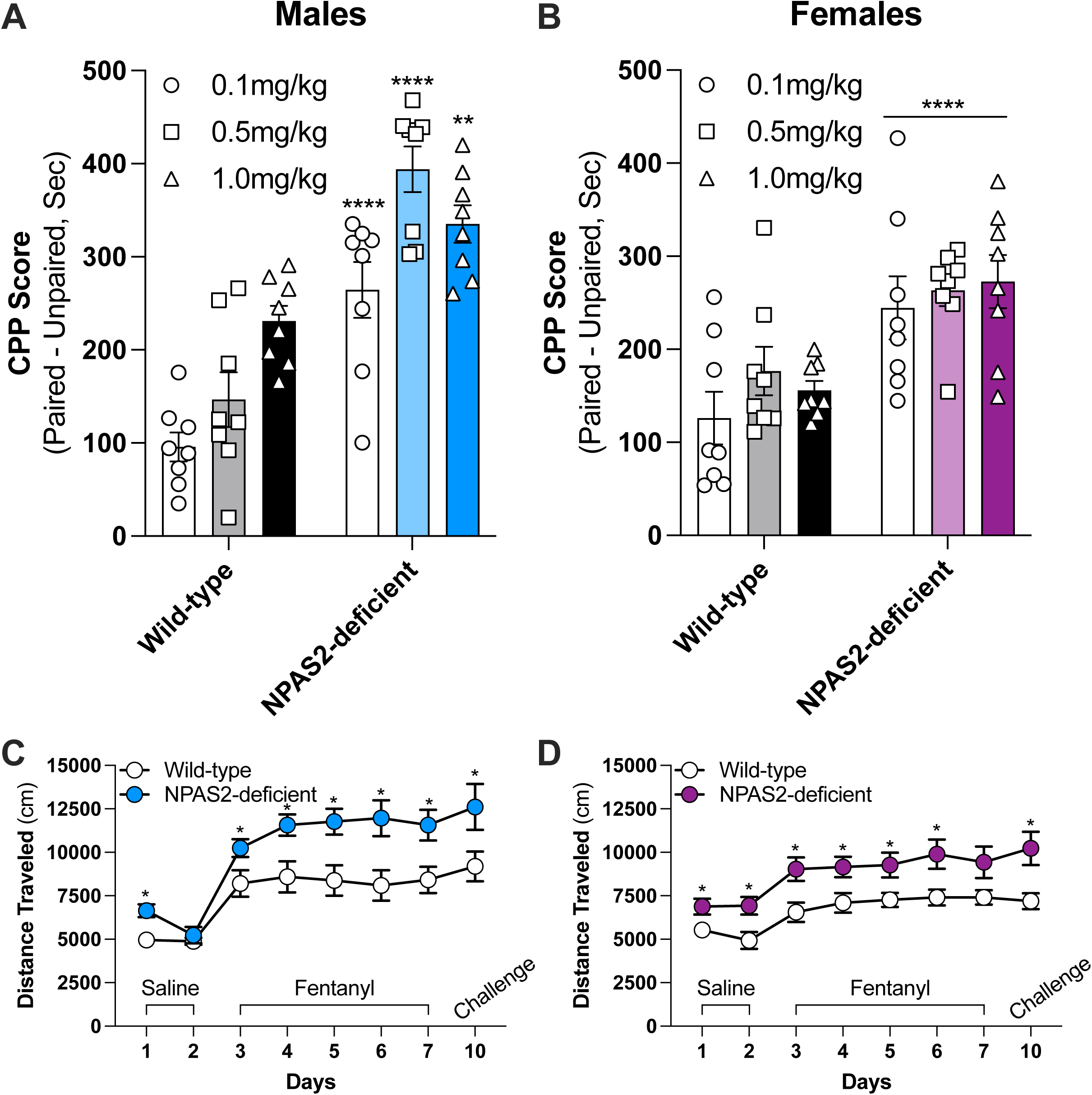
NPAS2 deficiency promotes fentanyl conditioned place preference and locomotor sensitization in both male and female mice. Conditioned place preference (CPP) across different doses of fentanyl (0.1, 0.5, and 1.0mg/kg) in **A)** male and **B)** female wild-type and NPAS2-deficient mice. Preference score is fentanyl paired deducted from the unpaired side in seconds. Locomotor as measured by distance traveled in centimeters in response to saline (days 1-2) or fentanyl (days 3-7), followed by a fentanyl challenge on day 10 in **C)** male and **D)** female wild-type and NPAS2-deficient mice. Data represented as mean ± SEM. *, p<0.05; **p<0.01; ****, p<0.0001.

### Sex-specific effects of Npas2 deficiency on operant oral self-administration of fentanyl

We trained mice to self-administer fentanyl by pressing an active lever to dispense a fentanyl-containing solution into an oral dipper. Mice learned to press the active lever and orally consume the fentanyl solution. NPAS2-deficient male mice lever-pressed to administer more rewards than wild-type mice across each of the session days (**Figure 2A**; main effects of genotype, F(1, 14) = 4.6, p=0.049, and day, F(5.36, 75), p<0.0001). No differences were found for correct and incorrect responses between wild-type and NPAS2-deficient male mice (p>0.05). Despite increased reward responding, the volume of fentanyl consumed during each session was similar between genotypes (**Figure 2C**). In female mice, NPAS2 deficiency led to consistently elevated administration of fentanyl rewards during each of the daily sessions (**Figure 2B**; main effects of genotype, F(1, 14) = 10, p=0.006, and day, F(4.1, 57.2), p=0.04). In contrast to males, NPAS2-deficient females consumed significantly more fentanyl solution per session than wild-type mice (**Figure 2D**; main effect of genotype, F(1, 14) = 8.8, p=0.01). There were no discernible differences between genotypes for male and female mice in active to inactive lever presses (**Figure 2E,F**) and to reach acquisition criteria (**Figure G,H**). Thus, male and female NPAS2-deficient mice displayed elevated responding for fentanyl reward, with only females consuming more fentanyl per session, resembling a possible sex-specific effect of sex on fentanyl-seeking behaviors.

**Figure 2.**
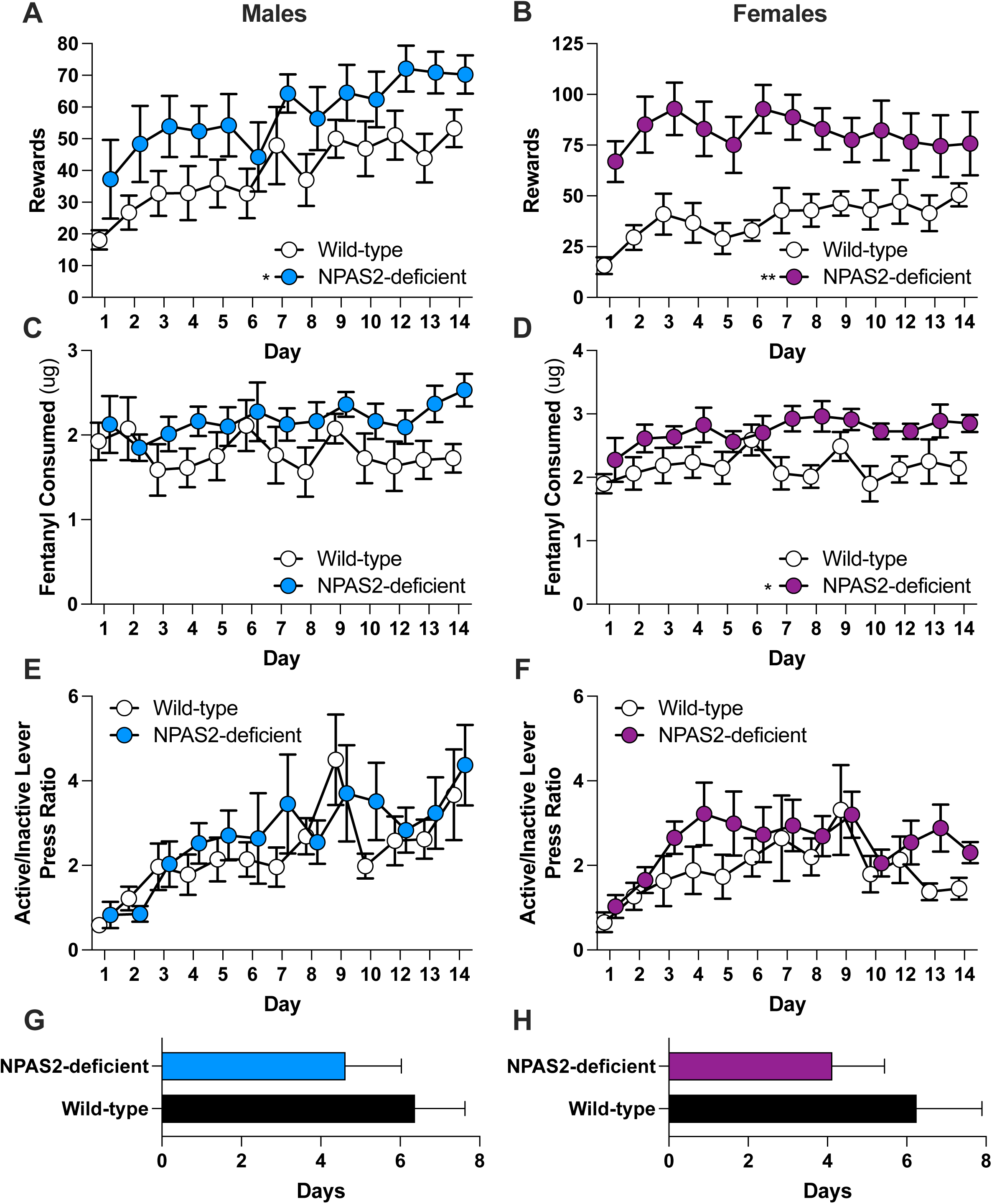
NPAS2 deficiency promotes fentanyl seeking behavior in both male and female mice. Mice were trained to press an active lever to dispense fentanyl containing solution into an automatic dipper to consume orally. Operant oral fentanyl self-administration behavior was conducted in wild-type and Npas2-deficient mice. Number of rewards self-administered by **A)** male and **B)** female mice. The amount of fentanyl consumed in ug per daily session by **A)** male and **B)** female mice. The ratio of active to inactive lever presses in **E)** male and **F)** female mice across daily sessions. The number of daily sessions to reach acquisition criteria for **G)** male and **H)** female mice. Data represented as mean ± SEM. *, p<0.05; **p<0.01.

Across different fentanyl doses, both wild-type and NPAS2-deficient males had similar reward responses (**Figure 3A**) and other measures including active to inactive lever presses, number of rewards self-administered, and progressive ratio values (**Figure 3B-D**). In males, both wild-type and NPAS2-deficient mice adjusted their behavioral responses as fentanyl dose increased (**Figure 3C**). In females, NPAS2deficiency led to significantly pronounced elevations of fentanyl rewards compared to wild-types across doses (**Figure 3G**; main effect of genotype, F(1, 70) = 40, p<0.0001), as reflected by changes in the active to inactive lever presses according to dose (**Figure 3H**). We also observed more rewards across doses in female NPAS2-deficient mice during progressive-ratio (**Figure 3I**; main effect of genotype, F(1, 42) = 18.2, p=0.0001). NPAS2-deficient females had higher motivation to self-administer fentanyl across doses (**Figure 3J**). Our findings suggest NPAS2-deficiency leads to higher motivation to self-administer fentanyl in female mice compared to males.

**Figure 3.**
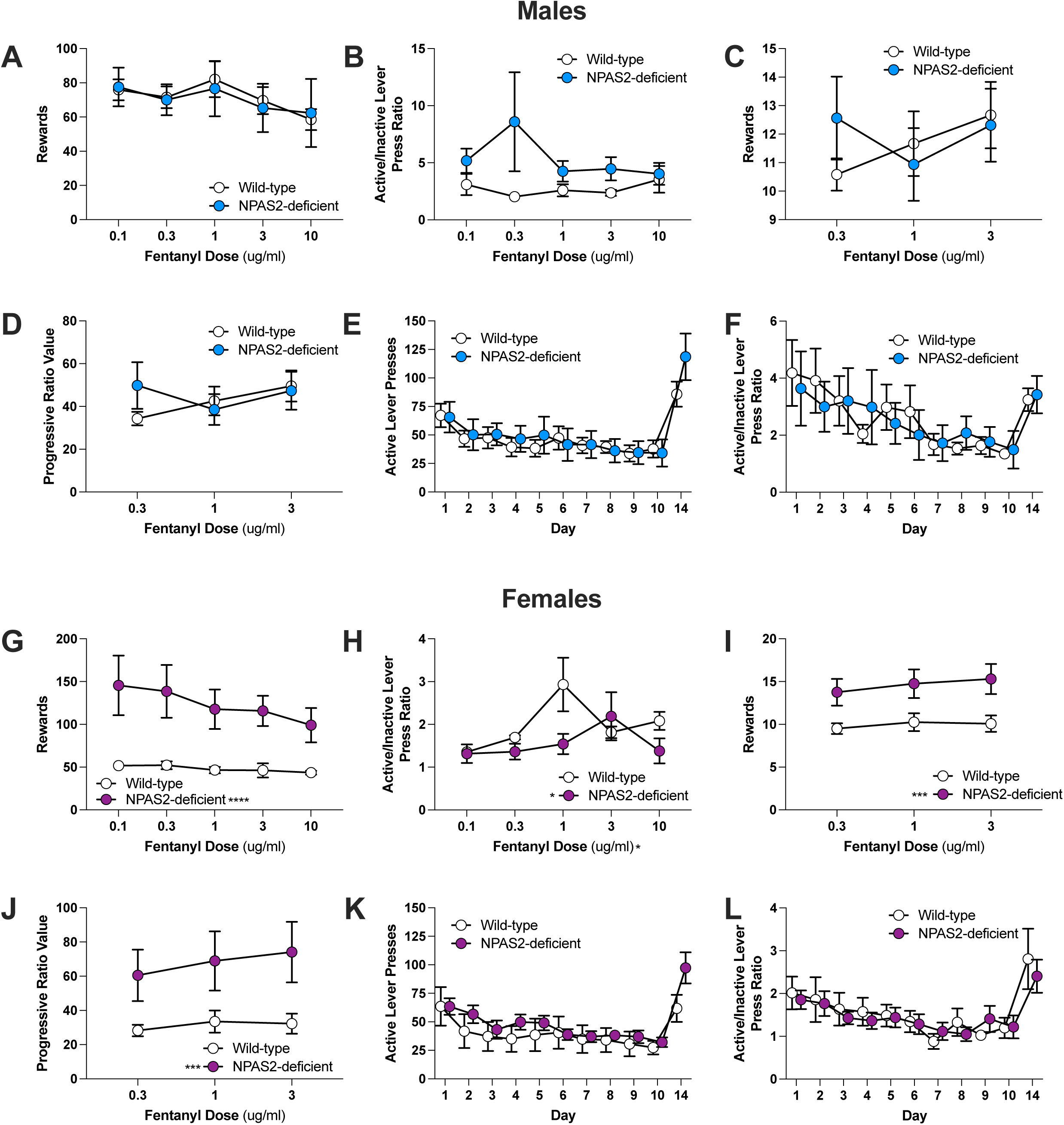
Sex-specific effects of NPAS2 deficiency on dose response and progressive ratio tasks during fentanyl self-administration behavior. **A)** Number of rewards self-administered and **B)** active to inactive lever presses during fentanyl dose response in male wild-type and NPAS2-deficient mice. **C)** Number of rewards self-administered, **D)** progressive ratio value (higher number of responses before receiving fentanyl dose, **E)** active lever presses, and **F)** active to inactive lever presses during progressive ratio task in male wild-type and NPAS2-deficient mice. **G)** Number of rewards self-administered, and **H)** active to inactive lever presses during fentanyl dose response in female wild-type and NPAS2-deficient mice. **I)** Number of rewards self-administered, **J)** progressive ratio value (higher number of responses before receiving fentanyl dose, **K)** active lever presses, and **L)** active to inactive lever presses during progressive ratio task in female wild-type and NPAS2-deficient mice. Data represented as mean ± SEM. *, p<0.05; **p<0.01; ***, p<0.001; ****, p<0.0001.

### Effects of NPAS2 deficiency on extinction and reinstatement of fentanyl self-administration behavior in male and female mice

Mice underwent extinction of fentanyl self-administration behavior over 10 successive sessions then re-introduced to cues (sound and light) previously associated with receiving a fentanyl reward following an active lever press. Both wild-type and NPAS2-deficient males reduced overall reward responding across each successive session (**Figure 3E,F**; main effect of day, F(10,154) = 6.1, p<0.0001). Female mice extinguished behavior across sessions similarly across genotypes (**Figure 3K,L**; main effect of day, F(10,154) = 4.1, p<0.0001). Further, both wild-type and NPAS2-deficient mice increased reward responding on day 14 when presented with cues associated with fentanyl (**Figure 3L**). Overall, no differences were found in the number of active to inactive presses during extinction and cue-induced reinstatement (**Figure 3F,M**).

### Cell type-specific role of Npas2 in striatal neurons on fentanyl reward-related behaviors

Since NPAS2 deficiency altered fentanyl reward-related behaviors, we aimed to assess whether fentanyl administration altered the expression of *Npas2* expression in the mouse nucleus accumbens. The nucleus accumbens had previously been shown to be an area of the striatum with dense, preferential enrichment of Npas2 expression and high levels of NPAS2-dependent transcription factor activity^6^. Other key circadian transcription factors, including the homologue, CLOCK, are also expressed differentially across the mouse striatum, and thus, may be influenced by opioids^7^. In a separate cohort of mice, we therefore measured the expression of *Clock* and *Npas2*. Notably, both acute and repeated administration of fentanyl reduced *Npas2* expression relative to saline in both males (**Figure 4A**; gene by condition interaction, F(2, 30) = 137.2, p<0.0001, with Sidak’s multiple comparisons test identifying both acute and repeated as significantly different than saline, p<0.0001) and females (**Figure 4A**; gene by condition interaction, F(2, 30) = 69.3, p<0.0001, with Sidak’s multiple comparisons test identifying both acute and repeated as significantly different than saline, p<0.0001) (**Figure 4A**). No changes in *Clock* expression were observed (**Figure 4A**; p>0.05).

**Figure 4.**
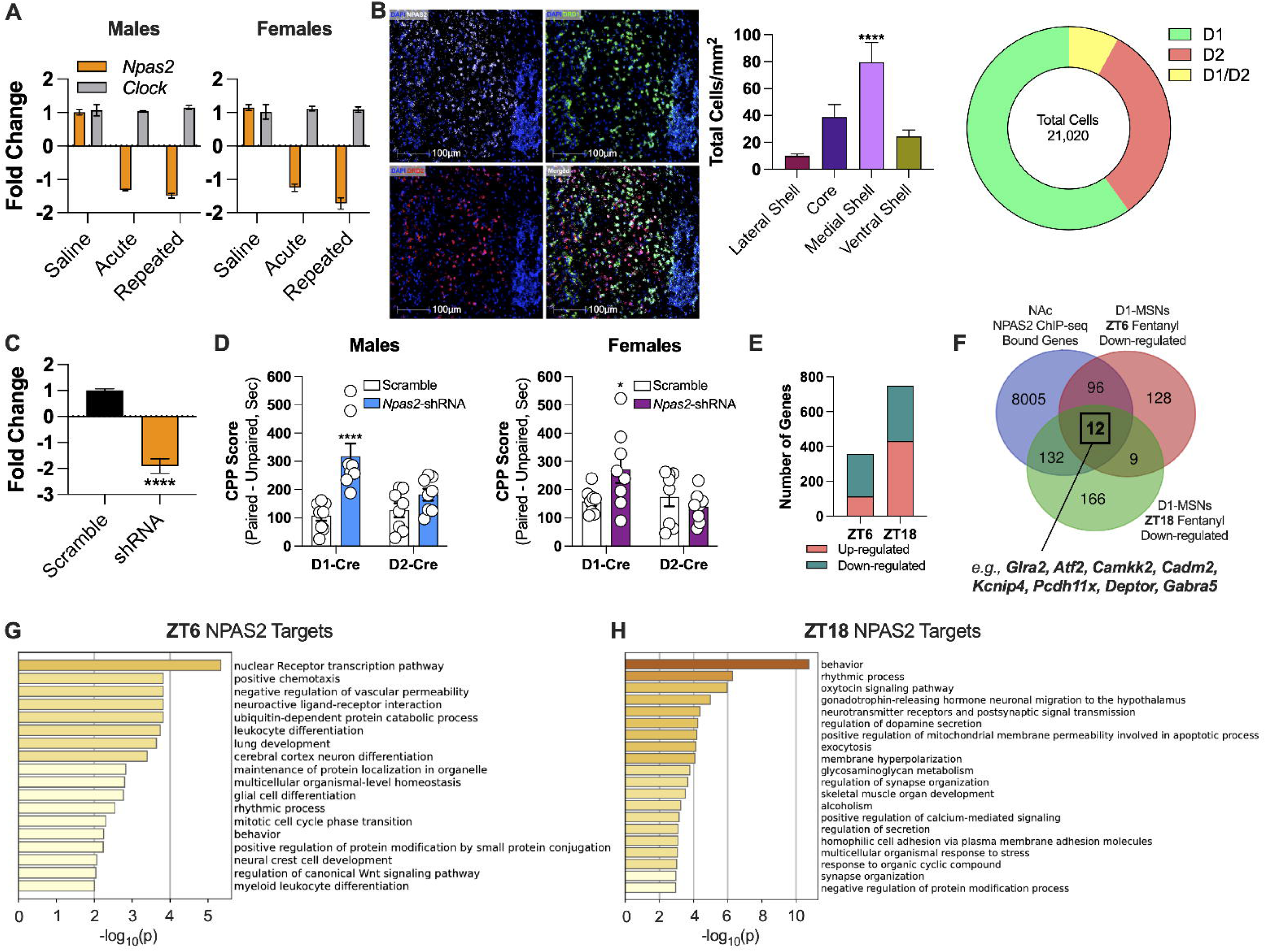
Cell type-specific knockdown of Npas2 in nucleus accumbens D1-MSNs leads to increased fentanyl conditioned place preference. **A)** Impact of acute or repeated fentanyl administration on *Npas2* and *Clock* expression in the nucleus accumbens of male and female mice. **B)** Labeling of *Npas2* with *Drd1* and *Drd2* in mouse nucleus accumbens. Preferential enrichment of *Npas2* expression in the medial shell of the nucleus accumbens and D1+ cells of the nucleus accumbens. **C)** shRNA-mediated knockdown of *Npas2* in D1-Cre mice. **D)** Fentanyl conditioned place preference (CPP) scores in male and female D1-Cre and D2-Cre mice. **E)** Number of genes up-regulated and down-regulated in D1-MSNs at ZT6 and ZT18 following repeated fentanyl administration in mice. **F)** Overlapping genes among Npas2-bound gene targets from Ozburn et al., 2015 and altered transcripts at ZT6 and ZT18 specifically in D1-MSNs of nucleus accumbens. Pathway enrichment of Npas2 gene targets and altered transcripts at **G)** ZT6 and **H)** ZT18. Data represented as mean ± SEM. *, p<0.05; ****, p<0.0001.

Using multiplex RNAscope fluorescent *in situ* hybridization, we mapped *Npas2* mRNA co-expression with both dopamine D_1_ (D1) and D_2_ (D2) receptor expression in medium spiny neurons of the mouse striatum. Prior work supported preferential expression of *Npas2* in D1+ striatal neurons relative to other cell types^7^. Consistent with prior work^7^, we found *Npas2* to be highly and preferentially expressed in D1+ striatal neurons compared to D2 and D1/D2 co-expressing neurons (**Figure 4B**). *Npas2* was highly expressed in the medial shell of the nucleus accumbens compared to other accumbal subregions within the ventral striatum (**Figure 4B**; F = 11, p<0.0001, medial shell versus lateral shell). No sex differences were found in *Npas2* expression across subregions of the nucleus accumbens (p>0.05).

To begin to investigate the cell type-specific of *Npas2* in fentanyl behaviors, we knocked down *Npas2* expression (**Figure 4C**) specifically in D1 or D2 striatal cells using Cre-dependent shRNA AAV in D1-Cre and D2-Cre male and female mice. In both males and females, cell type-specific knockdown of *Npas2* led to increased fentanyl CPP behaviors only in D1-Cre mice (**Figure 4D**; cell type by shRNA condition interaction in males, F(1, 28) = 3.9, p=0.01, with Sidak’s multiple comparisons test identifying increased CPP only in D1-Cre-specific *Npas2-*shRNA condition, p=0.0001; and cell type by shRNA condition interaction in females, F(1, 28) = 3.3, p=0.03, with Sidak’s multiple comparisons test identifying increased CPP only in D1-Cre-specific *Npas2-*shRNA condition, p=0.034). No effect of knockdown was observed in D2-Cre mice (**Figure 4D**; p>0.05). Together, our expression and manipulation studies identify a role for NPAS2 in regulating fentanyl reward-related behaviors via specific actions in D1+ striatal neurons.

### Involvement of NPAS2 in gene expression changes in D1+ striatal neurons by fentanyl administration

To identify genes that were impacted by fentanyl in D1+ striatal neurons, we conducted RNA-sequencing on RiboTag-labeled D1+ nucleus accumbens neurons from mice^12,21^ following the administration of fentanyl or saline. We collected D1+ cells at both ZT6 and ZT18, which represent the peak and trough of NPAS2-binding in the mouse nucleus accumbens, respectively^7^ (See Supplemental Tables 1 and 8 for Npas2 ChIP-seq genes). At ZT6, we observed the behavioral effects of Npas2 deficiency and knockdown. Most of the genes in D1+ cells altered by fentanyl were downregulated at both times of day (**Figure 4E**). Since NPAS2 is a circadian transcription factor, we directly compared downregulated genes to genes bound by NPAS2 in mouse nucleus accumbens across different times of day^7^. We identified 240 genes that were both downregulated in D1+ cells and bound by NPAS2 at ZT6 (108 genes) and ZT18 (144 genes) (12 genes common to both times of day; **Figure 4F**; Supplemental Tables 2-7, 9, and 10). Genes downregulated across both times of day which were also putative targets of NPAS2 included *Glra2, Atf2, Camkk2, Cadm2, Kcnip4, Pcdh11x, Deptor,* and *Gabra5* (**Figure 4F**). Several of these genes are key regulators of pathways we found to be enriched (**Figure 4G,F**). For example, *Glra2*^22^, *Atf2*^23^, *Camkk2*^24^, *Deptor*^25^, and *Gabra5*^26^ are involved in modulating circadian rhythms of neurons. Other pathways that were enriched included nuclear receptor transcription, chemotaxis, oxytocin signaling, and gonadotropic-releasing hormone signaling (**Figure 4G,H**). Together, our findings reveal putative targets of Npas2-dependent transcription specifically in D1+ neurons in nucleus accumbens that are altered by fentanyl.

### Loss of molecular rhythms in human nucleus accumbens identifies key gene network involving NPAS2

Finally, we translated our mouse findings to humans by investigating whether NPAS2 is involved in molecular rhythm alterations in opioid addiction. Specifically, we examined the loss of gene expression rhythms in the postmortem nucleus accumbens of individuals with OUD. Previously, we found 406 genes that lost rhythmicity in OUD^18,27^ (**Figure 5A**). We used ChEA3^19^, a web-server application developed to conduct transcription factor enrichment analysis, to predict transcription factors as regulators of the hundreds of genes that lost rhythmicity in OUD (**Figure 5A**), identifying many pathways (**Figure 5B**). NPAS2 was predicted to be among the top transcription factors (Supplemental Table 11). Notably, NPAS2 was connected to a larger network of genes that were altered in the nucleus accumbens of individuals with OUD (**Figure 5C**). NPAS2 is predicted to interact with several other transcription factors, including ARNTL (BMAL1) and RORB, a nuclear orphan receptor, both of which are known to dimerize with NPAS2 to regulate the molecular clock^3^. NPAS2 was also predicted to interact with NPAS3 and NPAS4, which are from separate families of proteins, known to be immediate early genes that are induced by stress and drugs of abuse^28,29^. Together, we report an association between NPAS2-dependent signaling in the nucleus accumbens of both humans and mice.

**Figure 5.**
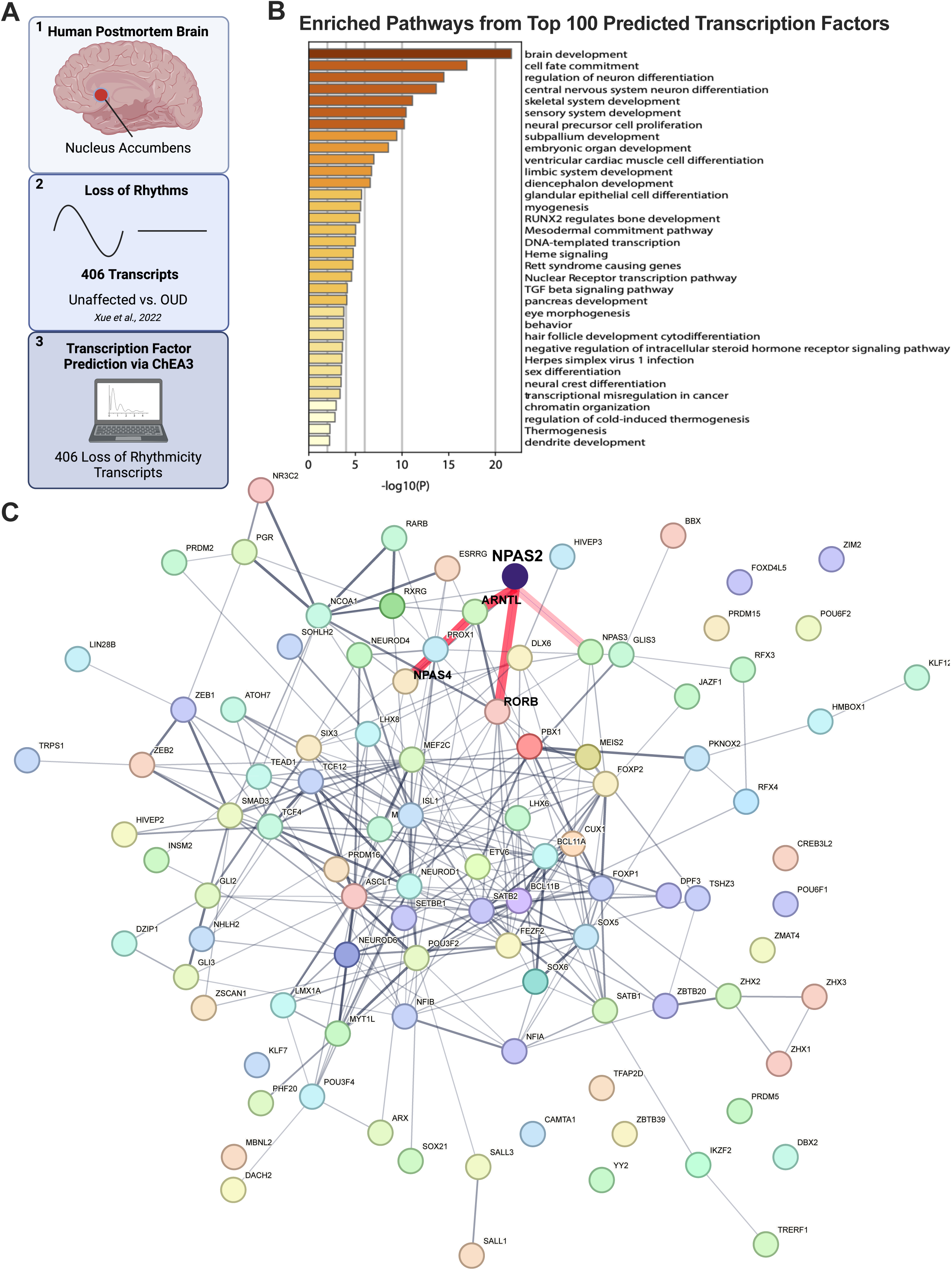
Prediction of transcription factors from loss of molecular rhythms in human postmortem nucleus accumbens in individuals with opioid use disorder. **A)** Schematic of workflow for predicting transcription factors based on transcripts that “lost rhythmicity” in human nucleus accumbens related to opioid addiction (see Xue et al., 2022). **B)** Pathways enriched from the top predicted transcription factors. **C)** Identification of NPAS2 in the transcription factor network predicted from transcripts that lost circadian rhythmicity in nucleus accumbens of individuals with opioid use disorder.

## DISCUSSION

The molecular and cellular mechanisms underlying the relationships between circadian rhythm disruptions and opioid addiction are largely unknown. Here, we report that chronic fentanyl administration leads to reduced expression of the circadian transcription factor, NPAS2, in the nucleus accumbens, a key brain region involved in opioid reward. Further, mice harboring a mutation in *Npas2*, which renders the transcription factor incapable of binding DNA to regulate transcription, exhibit elevated fentanyl conditioned reward and sensitization, and increased fentanyl self-administration behaviors. Fentanyl self-administration is particularly elevated in female NPAS2-deficient mice compared to males, suggesting sex-specific modulation of opioid-seeking behavior by NPAS2 actions. Female NPAS2-deficient mice also have augmented motivation for fentanyl across multiple doses. Interestingly, the expression of *Npas2* is enriched in the medial shell of the nucleus accumbens of both male and female mice, with preferential enrichment in D1 relative to D2 neurons^7^. Loss of NPAS2 function in D1 neurons leads to increased fentanyl conditioned reward, similar to our findings here in global NPAS2-deficient mice. Additional investigation identified several putative pathways that NPAS2 regulates specifically in D1-MSNs of nucleus accumbens, including nuclear receptor transcription, chemotaxis, vascular permeability, oxytocin, and gonadotropin-release hormone signaling, among others.

We previously reported sex differences in Npas2-dependent regulation of cocaine self-administration behavior^30,31^. Overall, female NPAS2-deficient mice tended to self-administer more cocaine with higher motivation and behaviors associated with craving than male mice, attributed, at least in part, to circulating ovarian hormones^31^. The caveat is that cocaine was self-administered intravenously, as here we used oral self-administration of fentanyl. Despite these differences, the response of *Npas2* in the striatum may be a key mechanism by which the brain responds to drugs of abuse and regulates subsequent motivation to seek and obtain the drug^30,32^. Differences between sexes in fentanyl self-administration behaviors is consistent with our prior work reporting NPAS2-deficient female mice displayed higher intensity and frequency of fentanyl withdrawal behaviors^33^, while males were hypersensitive to the development of fentanyl tolerance. We posit that hypersensitivity to fentanyl tolerance in NPAS2-deficient male mice dampen seeking behavior. In females, NPAS2-deficiency may intensify withdrawal symptoms^33^, driving augmented fentanyl-seeking behavior. Ongoing and future work will investigate the relationship between fentanyl self-administration and the development of physical dependence and tolerance among sexes in Npas2-deficient mice.

Prior work has also reported *Npas2* knockdown in D1-MSNs in the nucleus accumbens leads to reduced cocaine CPP^30^. We similarly found that *Npas2* knockdown in D1-MSNs leads to reduced fentanyl CPP, with no measurable effect in D2-MSNs, and subsequent reductions in expression following fentanyl administration in both sexes in both male and female mice. Reduced CPP in D1-MSN-specific NPAS2 knockdown was also similar to the impact of global Npas2-deficiency on fentanyl CPP and fentanyl-induced sensitization. Sensitization to repeated fentanyl administration by NPAS2-deficiency may be the result of alterations to excitatory synaptic strength and membrane excitability of post-synaptic MSNs in the nucleus accumbens^30^. Further work will need to investigate the direct role of NPAS2 in D1-MSNs and other MSN subtypes in response to fentanyl. Taken together, our findings strongly suggest that Npas2 is preferentially expressed in D1-MSNs and therefore more functionally relevant to opioid-related behaviors compared to other MSN subtypes.

Our findings also reveal the possible impact of sex on NPAS2 regulation of fentanyl reward, seeking, and craving behaviors in mice. Sex differences in circadian rhythms are attributable to, in part, circulating estradiol^34,35^. Indeed, ovariectomy of female mice slightly attenuated nightly elevations in cocaine self-administration behavior of NPAS2-deficient mice^31^. However, ovariectomy also increased the variability of cocaine-seeking behavior in female mice^31^, suggesting other sex-specific factors are involved in drug self-administration behaviors. Notably, locomotor sensitization and CPP were both increased in male and female NPAS2-deficient mice, while fentanyl seeking and taking behaviors were more consistently elevated in female NPAS2-deficient mice. Female NPAS2-deficient mice also consumed significantly more fentanyl during daily sessions relative to males. Despite the behavioral differences in NPAS2-deficient male and female mice, we surprisingly found no discernible differences in the impact of fentanyl administration on *Npas2* expression in the nucleus accumbens. This was also consistent with the effect of *Npas2* knockdown in D1-MSNs on fentanyl conditioned reward behaviors. Future studies are necessary to explicitly tease apart the impact of genetic and gonadal sex on NPAS2 actions in the nucleus accumbens and the sex-specific role of NPAS2 on opioid addiction.

Using prior NPAS2 ChIP-seq datasets^7^ and RiboTag pulldown of D1-MSNs, we identified overlaps in transcripts with known NPAS2 binding and fentanyl-induced alterations in D1-MSNs of mouse nucleus accumbens. Prior work demonstrates that DNA binding of NPAS2 in the nucleus accumbens resembles robust diurnal rhythmicity^7^, troughing at ZT6 and peaking at ZT18. We collected D1-MSNs from mice sacrificed at both ZT6 and ZT18, then examined the effects of fentanyl on cell type-specific transcriptomes. We focused primarily on downregulated transcripts since NPAS2 is a circadian transcription factor, acting in concert with other circadian proteins, to drive gene transcription at certain times of day. Interestingly, there were 12 transcripts that were downregulated at both ZT6 and ZT18 and known to be bound by NPAS2. Several of these transcripts regulate circadian rhythms or known to be regulated by molecular rhythms, including *Glra2, Atf2, Camkk2, Deptor,* and *Gabra5*. For example, *Atf2*^36^ and *Glra2*^37^ regulate neuronal activity rhythms, whereby reduced expression of these transcripts disrupts inhibitory signaling. Other transcripts include *Fmod*^38^ and *Pcdh11x*^39^, each of which have different roles in neuronal-synaptic signaling. Both *Cadm2*^40^, involved in synaptic adhesion, and *Kcnip4*^41,42^, a potassium channel, are associated with risk factors of substance use disorders, including neurodevelopmental disorders and increased risk-taking, impulsivity behaviors.

Our human brain studies demonstrated that the top predicted pathways from the overlap between NPAS2 bound transcripts and D1-MSN-specfic altered transcripts included nuclear receptor transcription and chemotaxis at ZT6 accompanied by behavior, rhythmic process, oxytocin, and gonadotropin-releasing hormone pathways at ZT18. *Nr3c1* is among the gene members of the nuclear receptor transcription pathway. NR3C1 is also known as the glucocorticoid receptor, involved in modulating the cellular response to circulating glucocorticoids^43^. *Rora* and *Rorb* are part of the nuclear receptor transcription pathway. RORs are nuclear orphan receptors with a myriad of roles in the brain^44^. RORs are also intimately involved in the regulation of molecular and cellular rhythms^3,44^. At ZT18, genes that regulate circadian rhythms are highly prevalent across enriched pathways such as *Egr1, Gabrb3, Ankfn1*, among others. *Egr1* is an immediate early gene involved in a number of processes associated with addiction^45–47^. Other notable transcripts include *Prkcg*, which is involved in modulating the functionality of mu-opioid receptors^48^, and an entire set of transcripts that regulate behavior and synaptic signaling—*Camk4*^49^, *Htr2a*^50^, *Syt2*^51^, *Kcnq3*^52^, and *Ntrk2*^53,54^. Collectively, our results support the direct action of NPAS2 to regulate a variety of processes involved in opioid actions that may be specific to D1-MSNs. Transcripts involved in a glucocorticoid signaling, synaptic plasticity, and circadian modulation of opioid actions on the brain may be involved in opioid reward and seeking behaviors.

Overall, our work provides new evidence that supports mechanisms associating circadian rhythm dysfunction with opioid addiction at molecular and cellular levels. Our prior work indicates Npas2 is involved in the development of tolerance to fentanyl and the severity of withdrawal symptoms in mice. We now extend these earlier studies to show that NPAS2 is a sexually dimorphic integrator of opioid actions that cuts across seemingly disparate phenotypes—tolerance, withdrawal, and reward to opioids. This is in stark contrast to the roles of other circadian genes in opioid actions and related behaviors^55^. Thus, modulating the action of NPAS2 and transcriptional targets may be a viable approach for developing therapeutics to prevent tolerance, mitigate withdrawal, and reduce opioid craving. Importantly, we predicted NPAS2 to be involved in a key signaling pathway of loss of rhythmicity in human postmortem nucleus accumbens in individuals with OUD, further supporting a key role for NPAS2 and linked transcript pathways in opioid addiction. Future studies will need to explore approaches to regulate NPAS2-dependent actions in the brain to treat behaviors associated with opioid addiction.

## Author Contributions

RWL, MLS, and ZF obtained funding for the project. RWL and ZF conceptualized and managed the project and provided supervision for data analyses and interpretations. KB, MAS, LMD, SK, SP, JOS, EF, and NMS developed and implemented self-administration experiments. JGK, SP, and ZF conducted the RNAscope multiplexed experiments, including analyses. SK, XZ, EF, and BW collected and processed mouse brain tissues for gene expression. LMD and SK completed cell type-specific RNA-sequencing assays. LMD, XX, and GCT completed RNA-sequencing analyses. PKP and RWL conducted stereotaxic viral surgeries, validation, and analyses. XZ and EF conducted conditioned place preference assays. KB, MAS, and RWL conducted the analyses, wrote the manuscript, and designed the figures. All authors participated in editing the manuscript for publication.

## Supplemental Information

**Supplemental Table 1 –** Genes Identified as Bound by NPAS2

**Supplemental Table 2 –** Differentially Expressed Genes Comparing Fentanyl to Saline in D1-MSNs

**Supplemental Table 3 –** Differentially Expressed Genes Comparing Fentanyl to Saline in D1-MSNs at ZT6

**Supplemental Table 4 –** Differentially Expressed Genes Comparing Fentanyl to Saline in D1-MSNs at ZT18

**Supplemental Table 5 –** Overlap of Differentially Expressed Genes in Fentanyl and NPAS2 Targets

**Supplemental Table 6 –** Overlap of Up-regulated Differentially Expressed Genes in Fentanyl and NPAS2 Targets

**Supplemental Table 7 –** Overlap of Down-regulated Differentially Expressed Genes in Fentanyl and NPAS2 Targets

**Supplemental Table 8 –** Genes with NPAS2 Bound to Promoter

**Supplemental Table 9 –** Genes with NPAS2 Bound and Down-regulated Differentially Expressed Genes in D1-MSNs

**Supplemental Table 10 –** Genes with NPAS2 Bound and Down-regulated Differentially Expressed Genes in D1-MSNs in ZT6 or ZT18

**Supplemental Table 11 –** CHEA Prediction Analyses of Transcription Factors in Human Nucleus Accumbens related to Opioid Addiction

